# Effect of osmoprotectants on the survival of bacterial endophytes in lyophilized beet roots

**DOI:** 10.1101/772939

**Authors:** Sonia Szymańska, Marcin Sikora, Katarzyna Hrynkiewicz, Jarosław Tyburski, Andrzej Tretyn, Marcin Gołębiewski

## Abstract

The increase of human population and associated increasing demand for agricultural products lead to soil over-exploitation. Biofertilizers based on lyophilized plant material containing living plant growth-promoting microorganisms (PGPM) could be an alternative to conventional fertilizers that fits into sustainable agricultural technologies ideas. We aimed to: (i) assess the diversity of endophytic bacteria in beet roots and (ii) determine the influence of osmoprotectants addition during lyophilization on bacterial density, viability, salt tolerance. Microbiome diversity was assessed based on 16S rRNA amplicons sequencing, bacterial density and salt tolerance was evaluated in cultures, while bacterial viability was calculated by using fluorescence microscopy and flow cytometry. Here we show that plant genotype shapes its endophytic microbiome diversity and determines physicochemical rhizosphere soil properties. Sea beet endophytic microbiome, consisting of genera characteristic for extreme environments, is more diverse and salt resistant than its crop relative. Supplementing osmoprotectants during root tissue lyophilization exerts a positive effect on bacterial community salt stress tolerance, viability and density. Trehalose improves these parameters more effectively than ectoine, moreover its use is economically advantageous.

## Introduction

Conventional agriculture practices negatively affect environment, e.g. by decreasing microbial diversity, soil quality, water supply and plant productivity (Gveroska et al., 2014; Baez-Rogelio et al., 2016). Wide adoption of sustainable agricultural technologies, e.g. biofertilizers, may significantly decrease the use of chemical fertilizers, reducing negative consequences of agriculture on the environment (Pretty, 2008; Malusá et al., 2011; Baez-Rogelio et al., 2016).

Biofertilizers are based on living plant growth-promoting microorganisms (arbuscular mycorrhizal fungi - AMF, plant growth-promoting rhizobacteria - PGPR, nitrogen fixing bacteria - NFB) and are key players in sustainable agriculture (Malusá et al., 2011). They can promote plant growth in several different ways (e.g. increasing availability of nutrients, synthesizing phytohormones or siderophores, fixing nitrogen), especially under unfavorable environmental conditions (e.g. drought, salinity) (Szymańska et al., 2016; Hrynkiewicz et al, 2019). Most of commercially available biofertilizers are based on two or more microbial beneficial strains, which is called ‘consortium’ (Malusá et al., 2011). Compared to single strains, consortia display increased spectrum of beneficial effect of inoculum on plants. However, criteria of strain selection are the crucial factor influencing the inoculum’s success and should be considered not only based on plant genotype compatibility but also environmental factors.

The methods of biofertilizers production, storage and application are diverse. Inoculation techniques are based on microorganisms application in liquid (sprays and drenches) or solid form (lyophilizates delivered to soil/growth substrate; Alori and Babalola, 2018). The most important problem in the preparation and storage technology of biofertilizers is maintaining high viability of microorganisms (Alori and Babalola, 2018). Lyophilization is well known and widely used technique extending viability (Prakash et al., 2013). To alleviate negative effect of low temperature on microorganisms in this technology several different stabilizers can be used e.g. nonreducing disaccharides, glicerol, skim milk (Park et al., 2002). Trehalose (α-D-glucopyranosyl-(1→1)-α-D-glucopyranoside) is a disaccharide present in almost all prokaryotic and eukaryotic organisms and exhibits high efficiency in protection of cells against low temperature, drying and osmotic stress (Park et al., 2002; Lee et al., 2018). Ectoine (1,4,5,6-tetrahydro-2-methyl-4-pyrimidinecarboxylic acid) is synthesized mostly by halotolerant and halophytic microorganisms and responsible for regulation of osmotic pressure in microbial cells, increasing their tolerance to osmotic stress (salinity) (Oren, 2008; Han et al., 2018 Roberts, 2005; Czech et al., 2018). Application of trehalose and ectoine in the process of lyophilization of endophytic microbiomes was tested in our work for the first time.

Biofertilizer’s efficiency analysed under laboratory conditions may not correspond to results obtained under field conditions (Baez-Rogelio et al., 2016; Parnell et al., 2016). This effect may be due to adverse effect of environmental conditions or autochthonic microorganisms on gene expression in microbial cells (Baez-Rogelio et al., 2016) or low competitiveness of microorganisms used as biofertilizers, i.e.they may be outgrown by autochthonic ones. This is why “plant microbiome” was proposed as new generation of inoculants (Compant et al., 2019). Inoculation of crops with microbiome and organic matter present in lyophilized plant roots seems to be a better solution to enrich microbial biodiversity of soil and crops with new endophytes.

Endophytes are bacteria and fungi that colonize the internal plants tissues without causing pathogenic symptoms (Hardoim et al., 2013) and can directly (nitrogen fixation, phosphate solubilization, siderophore and phytohormone synthesis) and/or indirectly (biocontrol agents) promote plant growth and development (Patle et al., 2018). Moreover, endophytes associated with halophytes possess high tolerance to salt stress (Abbas et al., 2018; Szymańska et al., 2018). Application of halotolerant microbes in sustainable agriculture e.g. in the increasing salinity tolerance of non-halophytic crops, is well known and was extensively studied (Yadav and Saxen, 2018; Etesami and Beattie, 2018; Szymańska et al., 2016, 2019).

Cultivated beets are one of crops whose direct ancestor (sea beet, *Beta vulgaris* ssp. *maritima*) still grows in the wild. This feature enables comparing traits in plants that are very close genetically (ca. 0.5% difference, Dohm et al. 2014), but whose ecology differs considerably. Moreover, as sea beet is a halophyte (Rozema et al. 2015), it seems to be a good candidate for a source of microorganisms that can be useful for crop beets improvement.

The goal of our study was twofold: i) to characterize root microbiomes of cultivated and wild beet and ii) to check if addition of osmoprotectants during lyophilization changed root bacterial community structure as well as microbiome salinity tolerance and viability. Specifically, we formulated five hypotheses: i) rhizospheric soils would differ in physicochemical parameters and bacterial community structure, ii) root endophytic communities would be different in the crop and its wild ancestor, iii) alpha diversity in the crop roots would be lower, iv) there would be more halophilic bacteria in sea beet roots, v) addition of osmoprotectancs would change bacterial viability and microbiome salinity tolerance.

## Materials and methods

### Experimental design

Sea beet (*Beta vulgaris* L. subsp. *maritima* L.) seeds were obtained from National Germplasm Resources Laboratory, Beltsville, MD, USA, while in the case of sugar beet (*B. vulgaris* subsp. *vulgaris* cv. ‘Huzar’) commercial seeds were bought from WHBC Poznań, Poland. Pot experiment was conducted from mid-March through mid-May 2017 in a greenhouse (Nicolaus Copernicus University in Toruń, Poland). Temperature was maintained at 22-24°C under natural lighting conditions. Healthy and uniform-sized seeds were placed in 5 l pots filled with 2,5 kg of garden soil (one seed per pot; 5 pots were prepared for each genotype, in total 10 pots were used). All plants for both investigated genotypes were arranged randomly on the green house benches. The plants were watered with tap water every two days, amount depended on the plants demand. After three months plants and rhizospheric soil samples were collected and analyzed as shown in Fig. 1.

**Figure 1.**
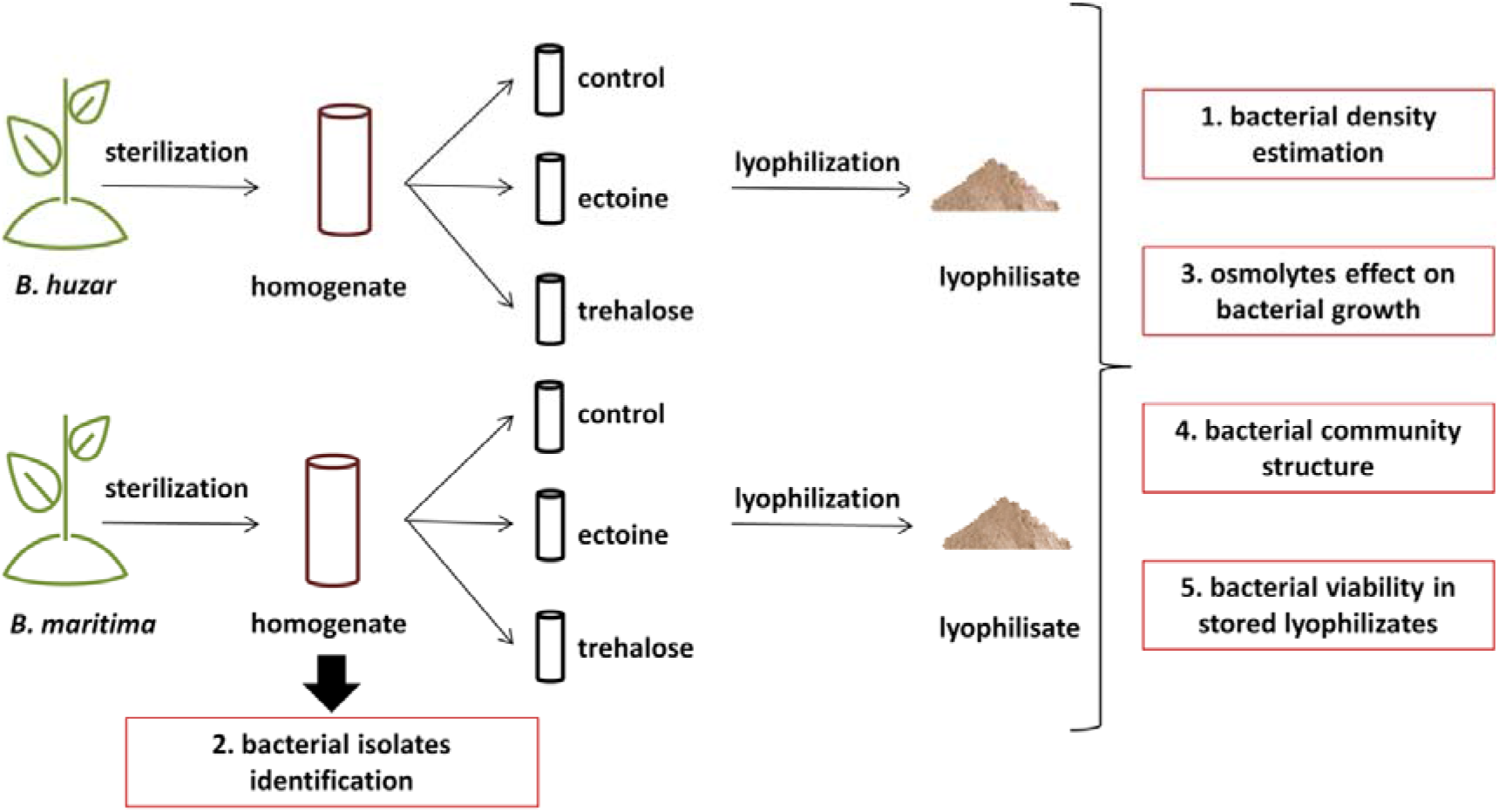
Experimental design.

### Soil analysis

Soil parameters (TOC, TN, CaCO_3_, P_citr_, pH, EC, Na, K, Ca, Mg, Cl, SO_4_^2−^) were analyzed as described earlier in Furtado et al., 2019.

### Plant and soil samples preparation

Plants were carefully uprooted, and 10 g of soil adhering to roots (rhizospheric soil) was collected, frozen at −80°C and lyophilized before DNA isolation for metagenomic analysis. Roots were washed with tap water to remove soil and were separated from shoots and leaves. Then, they were surface sterilized with 70% ethanol and 15% hydrogen peroxide mixture (1:1 v:v) for 5 min and subsequently rinsed six times with 0.9% NaCl. Efficiency of the sterilization process was evaluated by plating the last rinse on Luria-Bertani (Difco™ LB Agar, Miller) and potato dextrose extract (Lab A Neogen Company) media. Only properly sterilized plant material was used for subsequent analyzes. Approximately 100 g of fresh root material was homogenized in 100 ml of 0,9% NaCl by using surface sterilized (rinsed with 70% ethanol and UV-irradiated) blender. Homogenates were used to evaluate bacterial density and to prepare lyophilizates.

### Roots lyophilization

Homogenated sugar and sea beet roots were used to prepare three variants of lyophilizates including (1) control (without osmolytes addition) (2) trehalose and (3) ectoine supplemented. Three biological replicates were prepared for each tested plant species (9 samples per plant species, in total 18 samples were used for downstream analyzes). Either 1 ml of 0.9% NaCl (control) or xx g of trehalose (Tre) or yy g of ectoine (Ect) were mixed with 50 g of homogenized roots. The mixtures were lyophilized in (nazwa liofilizatora) until completely dry (approximately 24 h).

### Estimation of bacterial density

Serial dilutions were prepared directly from the homogenized fresh roots and lyophilizates resuspended in 0.9% NaCl (1:9 m:v). The dilutions (10^−3^ to 10^−8^) were plated in triplicates on LB plates supplemented with nystatin (Sigma, 100 μg/ml) to prevent fungal growth, and the plates were incubated for 5 days at 26°C. Colony counts (expressed as CFU per 1 g of fresh or dry weight for homogenates and lyophilizates, respectively) were based on plates with 30-300 colonies. At least six bacterial isolates were purified per experimental variant.

### Isolates identification by 16S rRNA gene sequencing

Genomic DNA was isolated from purified strains using GeneMatrix Bacterial and Yeast Genomic DNA Purification Kit (EurX) according to the manufacturer’s protocol with modified homogenization step (FastPrep®-24 bead-beater, one cycle of 20 seconds at 4.0 m/s). The DNA was analyzed spectrophotometrically (NanoDrop 2000). 16S rRNA gene fragment was amplified using 27F and 1492R primers (Daffonchio et al., 1998), following the procedure described in Szymańska et al. (2016). The products were purified with GeneMatrix PCR/DNA Clean-Up DNA Purification Kit (EurX) according to the manufacturer’s protocol. Sanger sequencing was performed with BrightDye Cycle Sequencing kit (Nimagen), using 40 ng of template DNA, 1.5 pmol of primer and 1 μl of kit and 1.5 μl of BD buffer in 10 μl volume. The reactions were EtOH/NaAc precipitated and read out at IBB PAS, Warsaw, Poland.

### Salt tolerance assessment

Salt tolerance of root bacterial communities was measured as OD_600_ after 5 days incubation at 26°C using 96-wells microtiter plate reader (Biolog Micro Station). 140 μl of LB medium supplemented NaCl to obtain final concentrations of 0, 50, 100, 150, 200, 300, 400, 500, 600, 700, 800, 900 mM were used per well. Inoculates were prepared by suspending 2 g of mortar-ground lyophilized roots in 18 ml of 0.9% NaCl and diluting the mixture ten times. The inoculates were filtered through 40 μm cell strainer (Biologix) to remove plant debris. Six test and two control wells were inoculated with 10 μl of filtered inoculate or 0.9% NaCl, respectively.

### Bacterial viability assessment: fluorescence microscopy and flow cytometry

Ten-miligram samples of ground lyophilizated roots were mixed with 10 ml of PBS (pH=7.4) and incubated for 2 days at 26°C with mixing. The mixtures were filtered through a 40 μm cell strainer (Biologix) and 2 ml were centrifuged for 3 min at 1000 × g at RT to pellet the residual plant debris. Cells in the supernatant were stained with Cell Viability kit (BectonDickinson) as per the manufacturer’s protocol, than bacterial viability was analyzed using fluorescence microscopy (after 6 and 12 months of storage) and flow cytometer (after 12 months storage). Preparations were photographed in red and green channel under 40 × magnification upon fluorescence excitation with 433 nm light on Axiostar plus fluorescence microscope (Zeiss) equipped with Delta Optical camera. Percentage of live cells was based on counts from at least 30 view fields per sample. Flow cytometric analysis was performed on samples stained as described above with FACS Aria III (BectonDickinson) using 488 nm laser for excitation. Fluorescence was collected at 530±30 nm (for thiazole orange - TO) and 616±26 nm (for propidium iodide - PI) bands and seventy-micrometer nozzle was used. Parameters were optimized basing on pure environmental strains and their mixtures analyses and autoclaved lyophilizates samples served as negative controls.

### Statistical analysis and bioinformatics

Bioinformatics analyses of Illumina reads was performed as described earlier (Thiem et al. 2018). Briefly, the reads were denoised, merged and chimeras were removed in dada2 (Callahan et al., 2016), then amplicon variant sequences were exported together with abundance information and processed in Mothur v.1.39 (Schloss et al., 2009): aligned against SILVA v.132 database, screened for those covering the 6428-22400 positions of the alignment, filtered to remove gap-only and terminal gap-containing positions, pre-clustered to remove residual noise and clustered into 0.03 dissimilarity OTUs. Representative OTU sequences were classified using naïve Bayesian classifier and SILVA database. Sanger reads were manually inspected in Chromas to remove obvious errors, the corrected sequences were merged with CAP3 (Huang and Madan, 1999), and classified using naïve Bayesian classifier (Wang et al., 2007).

Significance of differences between means was assessed with ANOVA test with Tukey’s post-hoc analysis implemented in Statistica 10.0 (StatSoft). Normality of data was tested with Shapiro-Wilk’s test and homogeneity of variance was assessed with Levene’s test. When the assumptions were violated non-parametric Kruskall-Wallis test was used.

## Results

### Rhizospheric soil physicochemical parameters are different for sugar and sea beet

Majority of tested parameters was higher in sugar beet soil, but only in cases of CaCO_3_ and Na^+^ the difference was statistically significant. On the other hand, OC, P, Ca^2+^, Mg^2+^ and N_t_ were higher in sea beet soil and for the latter the difference was significant (Table 1).

**Table 1.** Physico-chemical rhizospheric soil parameters (mean and standard deviation) obtained after three months of cultivation of sugar- and sea beet.

| Parameter\plant species | *B. huzar* | *B. maritima* |
| --- | --- | --- |
| OC (%) | 4.97 (1,646) | 5.66 (1,210) |
| N<sub>t</sub> (%) | 0.28 (0.032) | 0.34 (0.025) [↑] |
| CaCO<sub>3</sub> (%) | 1.88 (0.178) [↑] | 1.6 (0.107) |
| P<sub>cytr.</sub> [mg/kg] | 1183,08 (116,312) | 1253,29 (48.876) |
| pH | 7.1 (0.077) | 7.0 (0.074) |
| EC 1:5 [μS·cm<sup>-1</sup>] | 176,93 (57,826) | 142,28 (23,973) |
| Na<sup>+</sup> [mg·dm<sup>-3</sup>] | 16,80 (7,652) [↑] | 7,35 (1,885) |
| K<sup>+</sup> [mg·dm<sup>-3</sup>] | 2,95 (0,644) | 2,55 (1,111) |
| Ca<sup>2+</sup> [mg·dm<sup>-3</sup>] | 9,63 (2,291) | 12,93 (2,916) |
| Mg<sup>2+</sup> [mg·dm<sup>-3</sup>] | 1,28 (0,306) | 1,52 (0,223) |
| Cl<sup>-</sup> [mg·dm<sup>-3</sup>] | 46,02 (15,494) | 38,94 (2,890) |
| SO<sub>4</sub><sup>2-</sup> [mg·dm<sup>-3</sup>] | 39,35 (5,211) | 38,58 (7,998) |
| HCO<sub>3</sub><sup>-</sup> [mg·dm<sup>-3</sup>] | 94,55 (14,306) | 88,45 (29,063) |

[↑] significantly higher level based on Newman-Keuls test of rhizosphere soil parameter observed between the plant species.

### Bacterial diversity in sugar beet roots is lower than in its wild ancestor

Bacterial diversity, evenness and species richness were the highest in rhizospheric soil, regardless of the plant genotype. Lyophilized sea beet roots harbored more diverse community than sugar beet (Fig 2). The number of OTUs was ca. three times higher in the wild beet than in the crop (Fig. 2 AB), while the diversity was around 1.5 times higher (Fig. 2C), and evenness was ca. 1.3 times greater (Fig. 2D).

**Figure 2.**
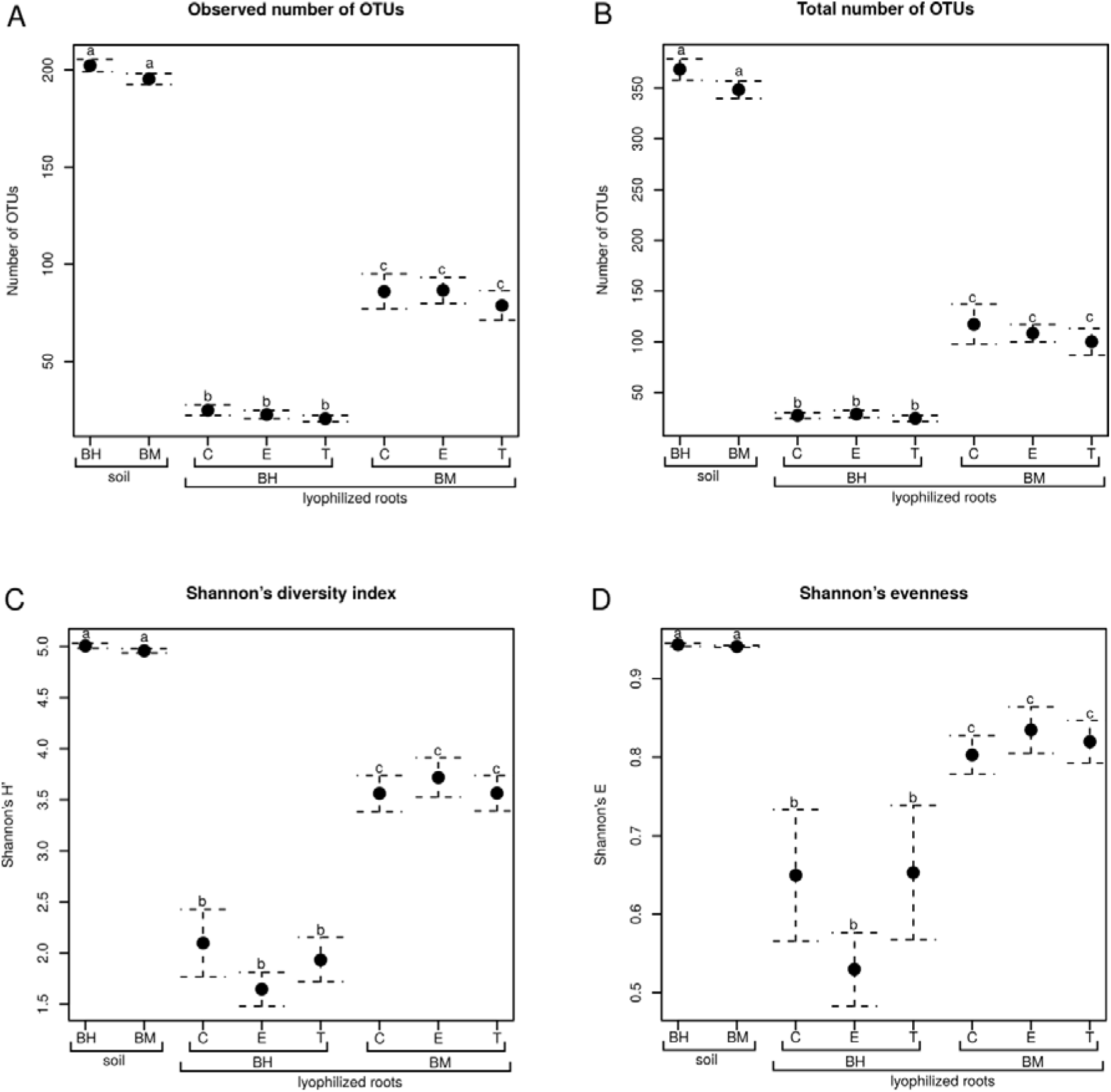
Species richness, evenness and diversity of bacterial communities in rhizospheric soils of sugar beet (Bh) and sea beet (Bm) and lyophilized roots of these plants untreated (C), and treated with ectoine (E) or trehalose (T). Means (n=8-32) are presented, whiskers show standard error of the mean (SEM), and significant differences (ANOVA, p<0.05) are denoted with different letters. Observed number of OTUs (A), estimated total number of OTUs (Chao1 index, B), Shannon’s diversity index (H’, C), Shannon’s evenness (D).

### Both endophytic and rhizospheric soil bacterial community is dominated by Proteobacteria

In total, 72 bacterial strains were identified, 35 coming from sugar beet and 37 from sea beet. Proteobacteria were the most frequent phylum in fresh roots of both sugar and sea beet, followed by Actinobacteria in the crop and Firmicutes in the wild plant. *Pseudomonas* and *Sphingomonas* were characteristic for fresh roots of sugar beet, while *Bosea* and *Sphingopyxis* were found exclusively in sea beet roots before lyophilization (Table 2).

**Table 2.**
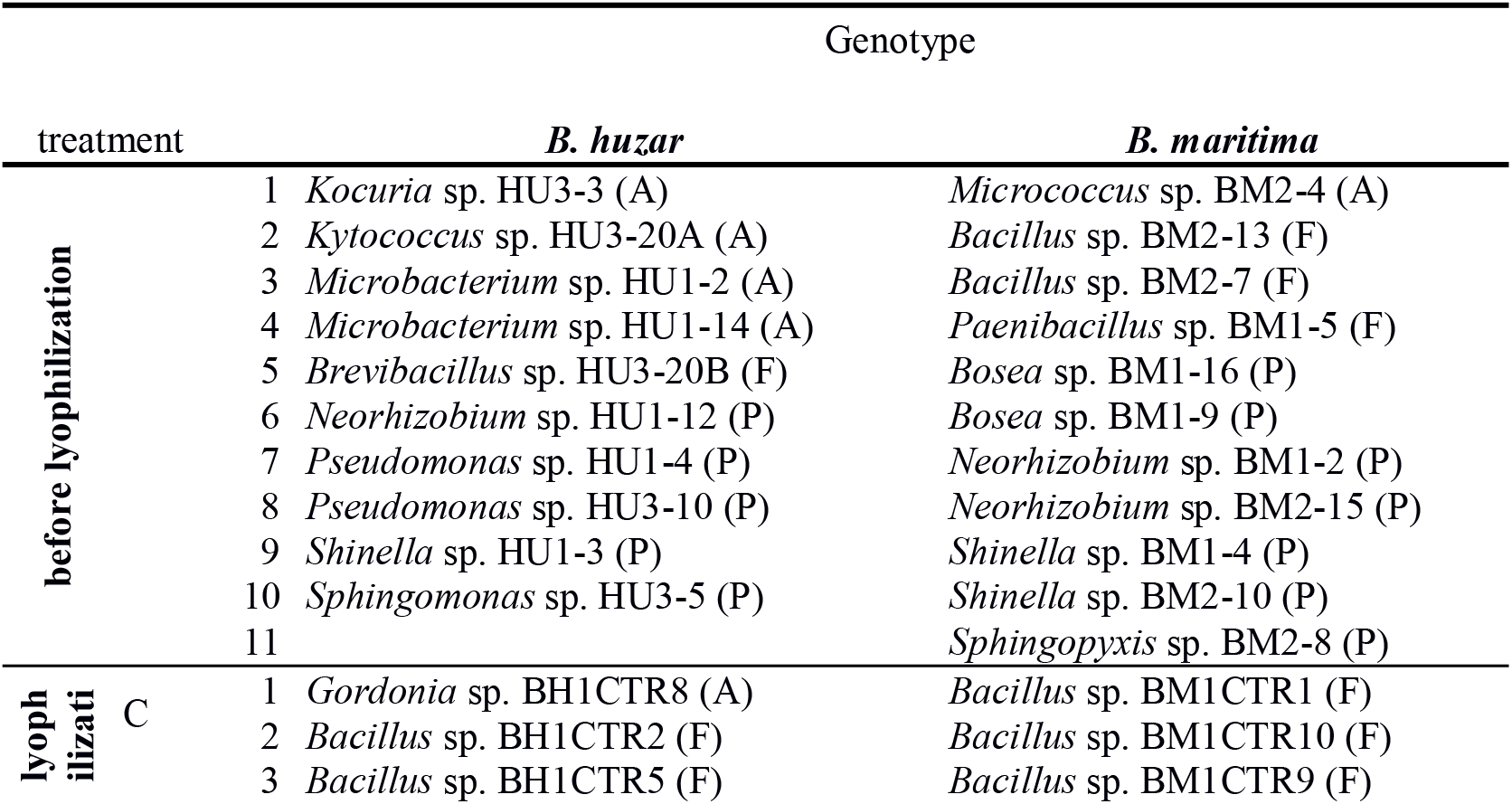
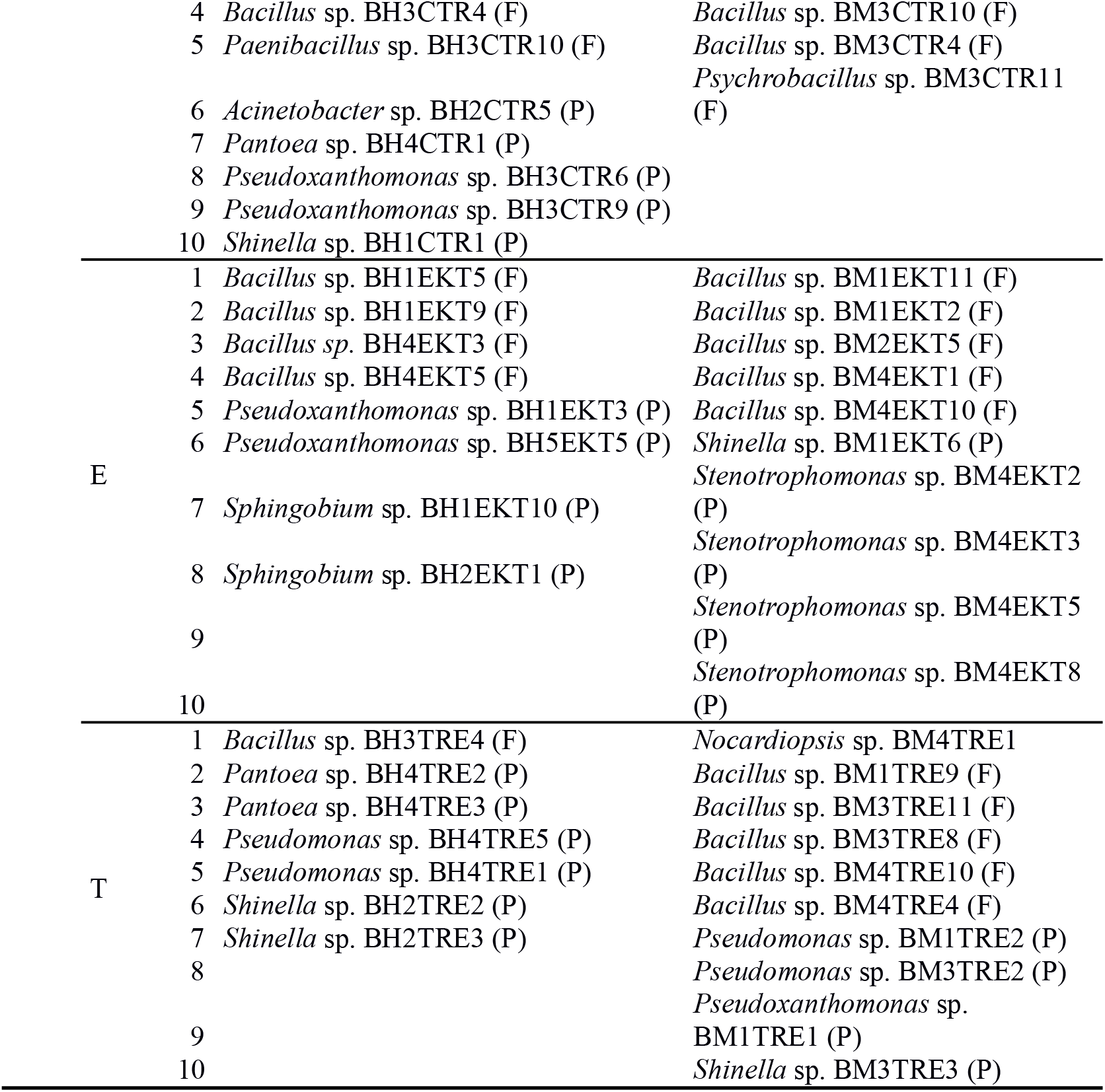
Identification of cultivable endophytic bacteria associated with roots of sugar- and sea beet before and after lyophilization without addition of any osmolyte (C) or supplemented either with ectoine (E) or trehalose (T).

There was no significant differences in taxonomic composition of rhizospheric soil bacterial communities of sugar- and sea beet at the level of phylum (Fig 3A). At the level of genus, three taxa were differentially represented, all of them belonging to Alphaproteobacteria: two Rhizobiales-belonging genera, *Pedomicrobium* and an unknown genus of JG34.KF.361 family as well as *Woodsholea* (Caulobacteraceae) were more abundant in the crop (fig. 3C). Differences in lyophilized roots communities were more pronounced, although still there were no taxa significantly differentially represented between osmolyte treatments. At the level of phyla Proteobacteria-derived reads were more abundant in libraries from sugar beet lyophilized roots, while Actinobacteria, Bacteroidetes, Acidobacteria, Verrucomicrobia and rare phyla were more abundant in its wild ancestor (Fig. 3B). Among genera significant differences were observed for *Stenotrophomonas* and *Bacillus* that were more abundant in the crop and for proteobacterial genera *Novosphingobium*, *Devosia* (Alphaproteobacteria), *Hydrogenophaga*, *Polaromonas* (Betaproteobacteria), *Rhizobacter* and *Tahibacter* (Gammaproteobacteria) as well as for rare and unclassified genera being more abundant in sea beet (Fig. 3D)

**Figure 3.**
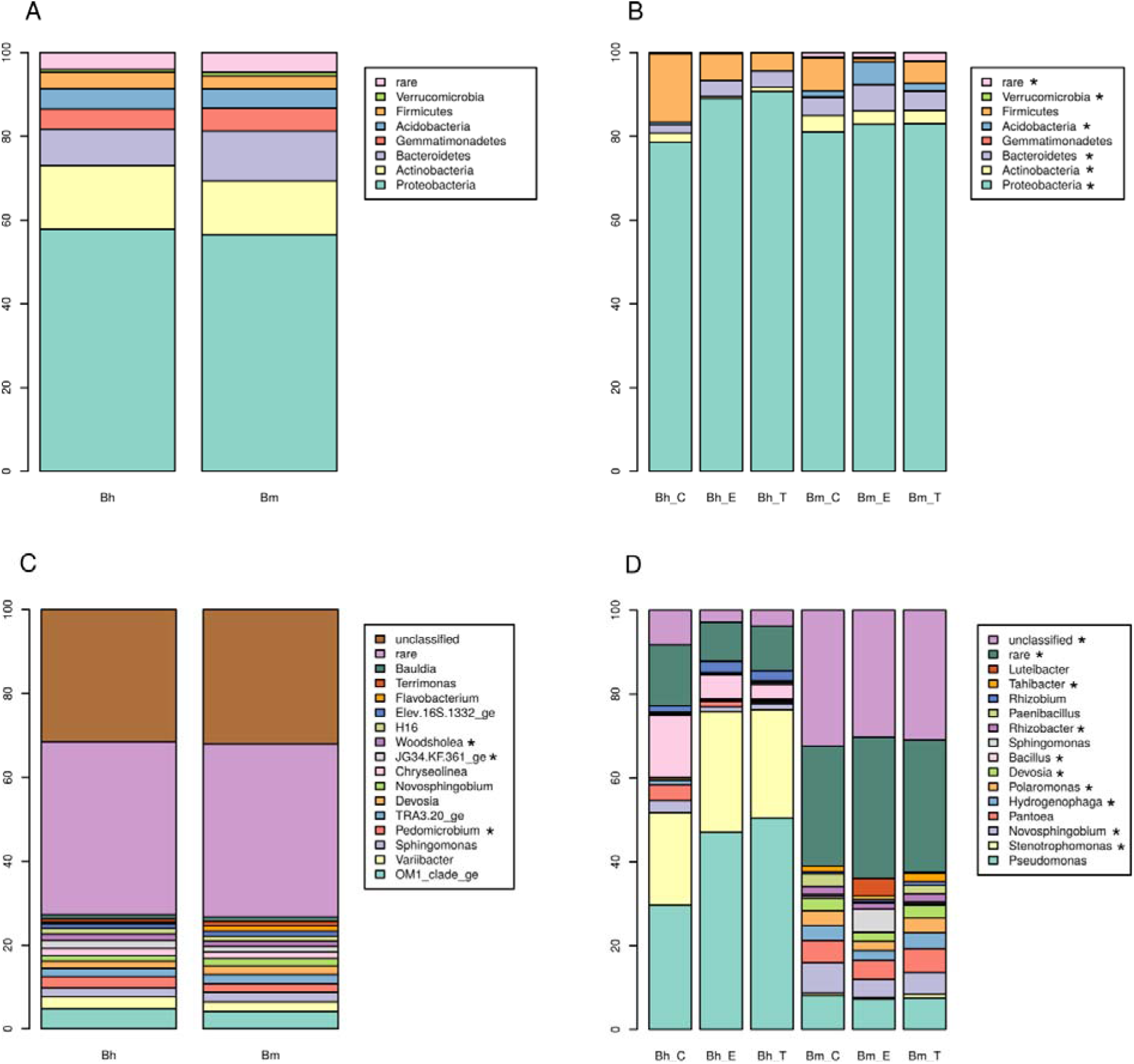
Taxonomic composition of bacteria communities in rhizospheric soils of sugar beet (Bh) and sea beet (Bm) (A, C) and lyophilized roots of these plants (B, D) untreated (Bh_C and Bm_C) and treated with ectoine (Bh_E, Bm_E) or trehalose (Bh_T, Bm_T) at the phylum (A, B) and genus (C, D) levels. Means (n=8-32) are presented, and significant differences between genotypes are marked with asterisks. No significant differences due to osmolytes were found.

### Culturable bacterial cell density in lyophilized roots depends on host genotype but not on osmolyte

Density of culturable root endophytic bacteria was higher in sugar beet lyophilizates than in sea beet (ANOVA, F=…, p<0.05, Fig. 4), regardless of the osmolytes addition. We observed no influence of osmolytes on sea beet endophytes density, while trehalose increased slightly, but significantly (ANOVA, F=…, p<0.05) the density in sugar beet samples (Fig. 4).

**Figure 4.**
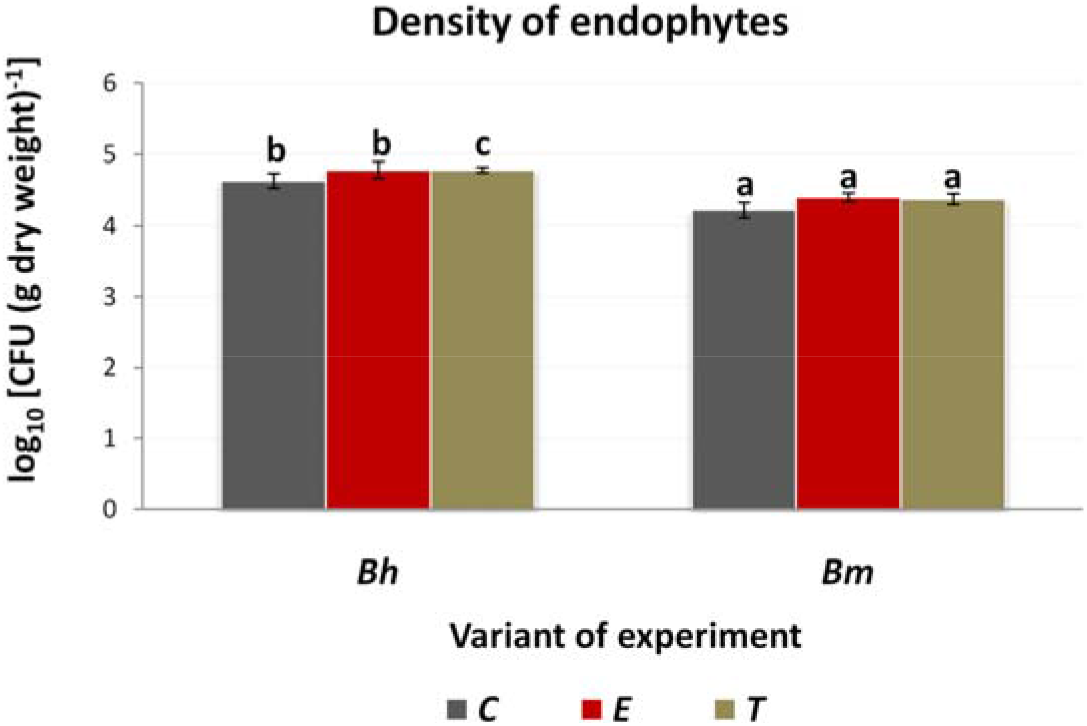
Density of endophytic bacteria (expressed as log_10_ CFU per g of dry weight) isolated from lyophilized sugar- and sea beet roots. Means (n = 3) ± standard deviation are presented. Significant differences between variants (ANOVA, p<0.05, with Tukey’s HSD; C – control untreated with any osmolyte, ectoine (E) or trehalose (T) treated) were marked with different letters.

### Sea beet endophytes are more salt tolerant than sugar beet ones

Increasing salinity negatively affected growth of culturable fraction of microbiome regardless of origin (sea- vs. sugar beet), however stronger effect was observed for *B. huzar*. In control treatment the growth was inhibited (final cell density below the critical level of 0.2 OD_600_) at 200 mM and 300 mM NaCl concentration for sugar and sea beet, respectively. Addition of osmolytes enhanced the growth in general and increased the inhibitory concentration to 400 and 700 mM, respectively (Supplementary Table 1). Influence of both osmolytes was similar with trehalose performing slightly better at high NaCl concentrations, and it was greater for sea beet, than for sugar beet (Fig. 5).

**Figure 5.**
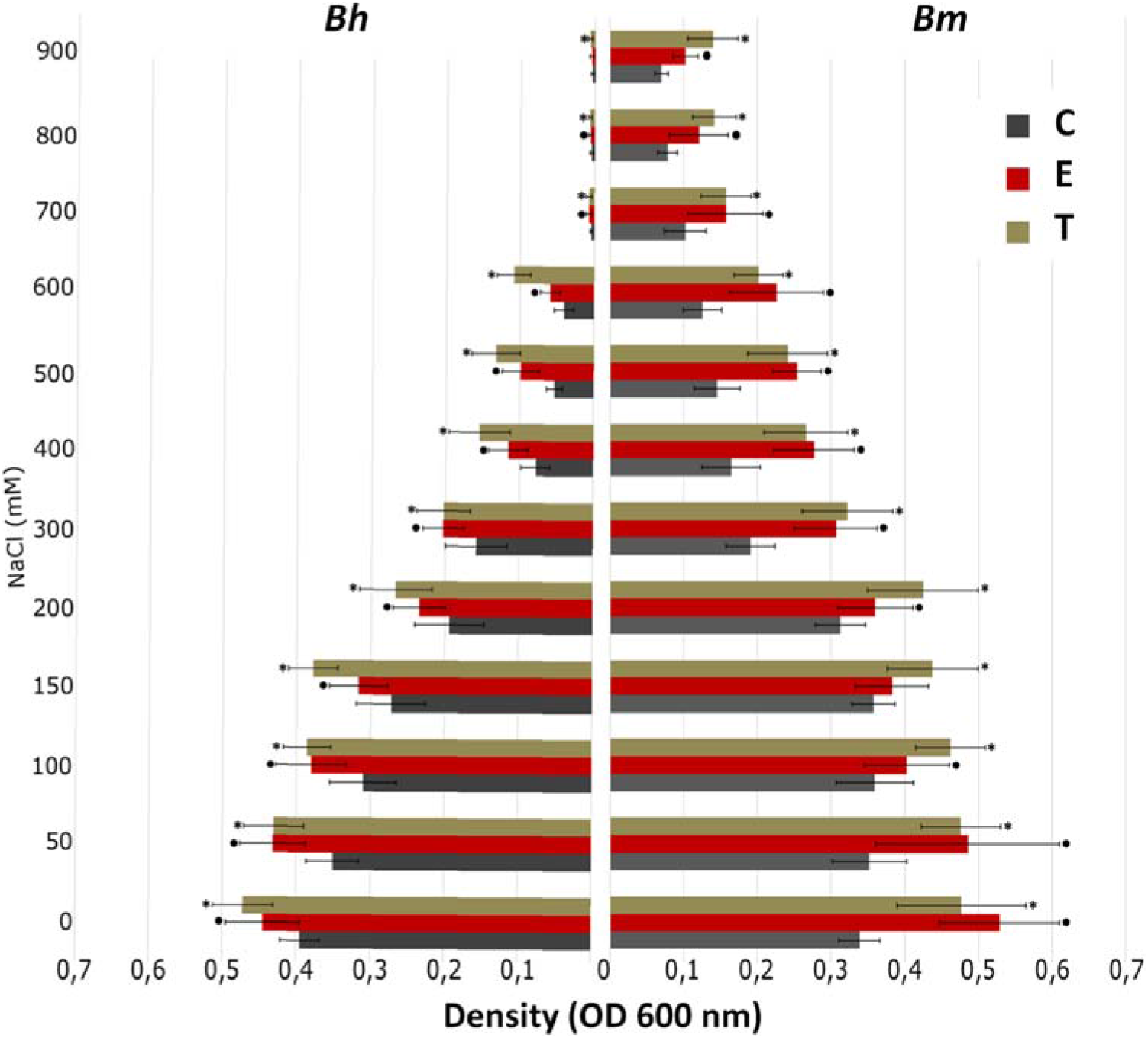
Osmolytes effect on growth of LB cultures inoculated with lyophilized sugar and sea beet roots untreated with any osmolyte (C), treated with ectoine (E) and trehalose (T). Means (n = 4-6) ± standard deviation are presented. Significant differences between treatments (ANOVA, p<0.05, with Tukey’s HSD) are marked with asterisks.

### Osmolytes enhance viability of Proteobacteria in lyophilized root samples

No influence of osmolytes on diversity (Shannon’s H’), observed, as well as estimated total (Chao1) OTU richness and Shannon’s evenness was observed in 16S rRNA gene libraries (Fig 2). Lyophilization without addition of osmolytes eliminated Actinobacteria and promoted Firmicutes, while addition of osmolytes (particularly trehalose) caused increase of Proteobacteria. (Table 2). To the contrary, community structure assessed via 16S rRNA gene fragment sequencing was similar, regardless of the osmolyte treatment (Fig. 2B,D).

### Bacterial viability in lyophilized roots were associated with plant genotype and osmolyte

Bacterial viability in lyophilized roots of sugar beet was consistently higher than in roots of its wild relative, regardles of osmolyte treatment, storage time and measurement methodology. Both osmolytes used increased the viability compared to control, regardless of genotype, but the effect of trehalose was more pronounced (Fig. 6).

**Figure 6.**
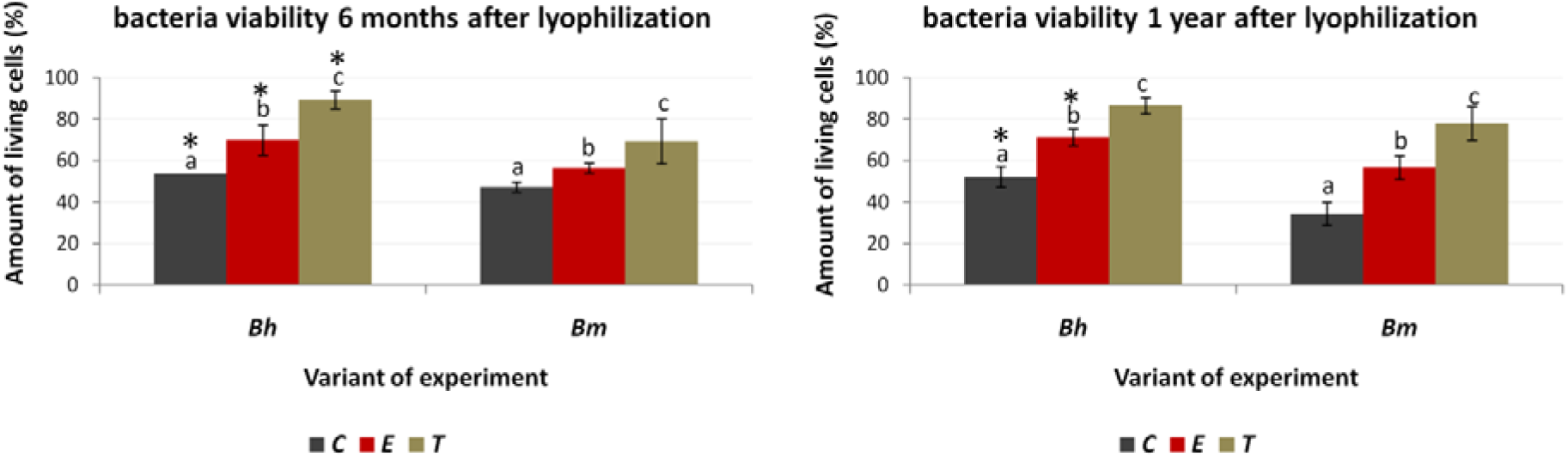
Bacterial viability in lyophilized beet roots. Viability measured with BD Cell Viability kit under fluorescence microscope (AB) and using flow cytometer (C). Means are presented and statistically significant differences are marked with differing letters.

## Discussion

### Bacterial diversity in beet rhizosphere

Exudates play pivotal role in shaping the rhizosphere ecosystem (Bashir et al., 2016). Differences in rhizospheric soil physicochemical properties observed in our study, may be due to greater nutritional demands of the plant (TN, Na) or varying exudates composition (OC), as it was found that rhizodeposition is the primary organic carbon source in rhizosphere (Bashir et al., 2016). Alternatively, they might be caused by changes in microbial activity resulting from exudation or interaction between microorganisms (Nannipieri et al., 2008; Shi et al., 2011).

Greater microbiome diversity in rhizosphere compared to endosphere was commonly observed, and resulted from natural plant selection mechanisms (Kandel et al., 2017; Liu et al., 2017; Cheng et al., 2019). Accordingly, in our study, the higher bacterial diversity, evenness and species richness were noted for rhizosphere soil of both investigated genotypes, than for roots. At the same time, in spite of slightly different TN, OC and Na levels, microbiome composition and diversity were similar in rhizospheric soils of both studied beets. This observation could be explained by the use of the same garden soil and short culture period, not allowing the differences to fully manifest. Culture-independent analysis revealed that dominating bacterial phyla were the same as those observed in rhizosphere of many plant species e.g. barley, alfalfa, wheat (Velázquez-Sepúlveda et al., 2012; Bulgarelli et al., 2015; Kumar et al., 2019). Only a few differences were noted at the genus level, mainly concerning Alphaproteobacteria. *Pedomicrobium* as well as JG34.KF.361_ge, more frequent in sugar beet, represent *Rhizobiales,* an order known for organisms that establish beneficial interactions with plants and comprises numerous bacteria with nitrogen-fixing capability (Erlacher et al., 2015). The observed lower TN level in the sugar beet rhizosphere may indicate crop’s higher demand for nitrogen. Tsurumaru et al. indicated that *Mesorhizobium* and *Bradyrhizobium,* also belonging to *Rhizobiales,* play an important ecological role in the taproot of sugar beet (2015). Moreover, Abdel-Motagally and Attia showed that higher levels of nitrogen (N) and potassium (K) significantly affect the growth parameters of sugar beet (2009). Both elements were generally recognized as crucial for obtaining higher yields of this crop, favorably affecting organic metabolites biosynthesis and improving nutritional status (Abdel-Motagally and Attia, 2009).

### Bacterial diversity in beet roots

The higher diversity both in rhizo- and endosphere of the wild plant compared to its crop counterpart was observed (Zachow et al. 2014; Ofek-Lalzar et al., 2016). It was hypothesized that beneficial endophytes associated with wild plants were absent or fewer in domesticated crops (Ofek-Lalzar et al., 2016). Sugar beet as a cultivated plant grows under more controlled conditions regulated by farmers, while sea beet grows mainly in highly saline and nutrients poor coastal soil (Tan et al., 2017). Growth under adverse environmental conditions requires support of microorganisms with a wide range of beneficial metabolic properties tailored for specific plant needs (Szymańska et al., 2018). The loss of high tolerance to salt stress during the process of sea beet domestication was demonstrated (Rozema et al., 2014) and might be associated with the loss of microbes that increased tolerance of this plant to salinity. Concordantly, despite the lack of differences in rhizospheric soil microbial composition, lower diversity of endophytes in sugar beet compared to its wild ancestor was noted in our study. This difference might be explained by varying root system architecture, with fibrous root system of sea beet providing more opportunities for bacteria to enter the endosphere (Kandel et al., 2017), which affects stochastic community assembly. On the other hand microbe selection can be driven by the genetic makeup of two studied subspecies. We observed that sea beet caused decrease in the soil Na level, suggesting aaccumulation of Na ions in wild plant tissues. Accordingly, there was an increase in community salinity resistance in this plant, which pointed at higher level of halotolerant and halophytic microorganisms.

In general, endophytic microbiome diversity and composition is related to soil properties as well as plant ecology and physiology (Miliute et al., 2015). Members of only three phyla (Proteobacteria, Actinobacteria and Firmicutes) were cultured in our experiment, this may be related to their high ability to grow on commercially available media (Milute et al., 2015; Szymańska et al. 2016a, Szymańska et al. 2016b; Brígido et al., 2019). Miliute et al. (2015) emphasized that Proteobacteria distinctly predominate among culturable plant endophytes, then the presence of Firmicutes and Actinobacteria is common, and Bacteroidetes occur slightly less frequently.

16S rRNA libraries generated in our study were dominated by the four phyla (Proteobacteria, Actinobacteria, Firmicutes, Bacteroidetes) commonly found in endosphere of glycophytes including maize (*Zea mays* L.), *Dactylis glomerata* L., *Festuca rubra* L. and *Lolium perenne* L. as well as in halophytes such as *Salicornia europaea* or para grass (*Urochloa mutica*) (Mukhtar et al., 2016; Zhao et al., 2016; Wemheuer et al., 2017; Correa-Galeote et al., 2018; Szymańska et al., 2018). Sea beet was characterized by significantly higher frequency of Actinobacteria, Bacteroidetes, Acidobacteria, Verrucomicrobia and rare phyla compared to sugar beet, where Proteobacteria were observed more often. Zachow et al. (2014) noted more Actinobacteria, Bacteroidetes and Verrucomicrobia in rhizosphere of wild beet cultivated in coastal soil than in sugar beet. This fact, together with our results, may point at these bacterial taxa being preferred by sea beet regardless of soil.

Our 16S rRNA gene sequencing results also revealed significantly higher abundance of certain genera in sea beet endosphere, including: *Novosphingobium*, *Devosia* (Alphaproteobacteria), *Hydrogenophaga*, *Polaromonas* (Betaproteobacteria), *Rhizobacter* and *Tahibacter* (Gammaproteobacteria) as well as certain rare and unclassified bacteria. These microorganisms comprise extremophiles, e.g. Polaromonas (Margesin et al., 2012; Gawor et al., 2018) or Hydrogenophaga (Khan and Goel, 2008) and organisms modulating plant stress response, such as Novosphingobium (Etesami and Beattie, 2018; Vives-Peris et al., 2018). In our study, only *Stenotrophomonas* and *Bacillus* species were more frequent in roots of sugar beet than of sea beet. *Stenotrophomonas* and *Pseudomonas* sp. were identified in rhizospheric soil of sugar and sea beet while first of them and *Staphylococcus* sp. were mainly observed for crop rhizosphere. Sea beet microbiome was found to be more diverse than that of sugar beet (Zachow et al., 2014), which explains greater number of rare taxa. It was found that sugar beet rhizosphere was more frequently colonized by strains with antagonistic activity against plant pathogens and/or stress protection activity, while abiotic stress-releasing ones were more often found in sea beet’s rhizosphere (Zachow et al., 2014). These facts together with our results suggest that pre-adaptation to stress observed in sea beet transcriptome (Skorupa et al., 2019) may also take place at the level of microbiome serving as a helper.

### Osmoprotectants enhance bacterial viability and diversity in lyophilized beet roots

Significantly higher cell density of culturable bacteria observed in sugar beet lyophilized roots can be attributed to high content of sucrose (Haankuku et al., 2015). This sugar acts as a natural osmoprotectant, allowing a better viability of microorganisms during lyophilization (Bircher et al., 2018). Another explanation of obtained results can be associated with higher ability of sugar beet endophytes to grow on solid medium.

Sea beet endophytic microbiome was found to be more resistant to salinity. Microorganisms present in a more saline sea beet tissue most likely developed mechanisms of adaptation to high salt level, which provided them ability to grow in higher NaCl concentrations compared to the sugar beet microbiome. This fact may be related to higher sodium accumulation in this plant tissues (Skorupa et al. 2019), which caused drop in soil sodium concentration observed in our study.

Salinity-induced changes in community structure and adverse effects on microbial density, activity, biomass were reported by many scientists (Yan et al., 2015, Zhang et al., 2019). The decrease in number of culturable microorganisms related to increasing NaCl concentration was noted even in the case of endophytes associated with halophytes (*Aster tripolium*, *Salicornia europaea*) (Szymańska et al., 2013; Szymańska et al., 2016). Obtained results were in line with the above trend, but apart from negative effect of salinity on sugar and sea beet bacterial density, a beneficial impact of trehaloze and ectoine on salt stress mitigation was demonstrated. Although ectoine is a major osmolyte in aerobic chemoheterotrophic bacteria and is considered as a marker for halophytic bacteria (Roberts, 2005), a slightly better effect of trehalose, was confirmed by the results of microscopic analyzes, flow cytometry and culture tests. Better performance of trehalose can be connected with its higher concentration used, which better counteracts the external osmotic pressure. Protective effect of trehalose is explained by “water replacement hypothesis” that states that the compound lowers the phase transition temperature of membrane phospholipids, by replacement of water molecules occurring around the lipid head groups (Berninger et al., 2017), thus protecting membrane structure (Nounjan and Theerakulpisut, 2012). The suggest that the use of trehalose is a better and more economic solution providing high viability of bacterial cells after lyophilization. In the case of sugar beet the above mentioned positive sucrose impact was enhanced by trehaloze addition. Similar effect was observed by Pereira et al. (2012) for rhizobial strains, where trehaloze worked better than sucrose/peptone mixture. In general, 16S rRNA gene sequencing results considering diversity of endophytes associated with sea and sugar beet root did not show any effect of applied osmoprotectants neither on alpha nor beta diversity of bacteria. This observation can be explained by the presence of ‘relic DNA’, i.e. DNA coming from non-viable cells (Carini et al., 2017) in lyophilized samples.

*Bacillus* sp. was the only species identified among the strains representing the Firmicutes phylum isolated from the lyophilized osmolytes treated roots of both investigated genotypes, in the control variant the presence of *Psychrobacillus* sp. and *Paenibacillus* sp. inside sea and sugar beet root was additionally found, respectively. The presence of the above-mentioned strains in the roots of the tested plants after lyophilization was associated with the commonly known their ability to endospore-forming and higher tolerance to environmental changes (Nicholson et al., 2000; Pham et al., 2015; Sáez-Nieto et al., 2017). Actinobacteria proved to be a very sensitive to lyophilization. Proteobacteria remarkably well tolerated the lyophilization, additional osmolytes promoted the incidence of culturable bacteria belonging to this phylum.

## Conclusions

Our research revealed that plant genotype played a pivotal role in the shaping of its endophytic microbiome diversity and physicochemical rhizosphere soil properties, mainly on salinity, but not soil bacterial community structure. Bacterial diversity was lower in sugar beet roots than in its wild ancestor tissues. At the same time sea beet endophytic microbiome was more salt resistant and consisted of genera characteristic for extreme environments.

Supplementing osmoprotectants during root tissue lyophilization had a positive effect on bacterial salt stress tolerance, viability and density. Trehalose proved to improve these parameters more effectively than ectoine, moreover its use was economically advantageous.

## Authors contribution

SS – performed experiments, analyzed data, drafted the manuscript, MS – performed experiments, helped in drafting the manuscript, KH – conceptualized the study, participated in writing the manuscript, JT – participated in writing the manuscript, AT – participated in writing the manuscript, MG – supervised the project, conceptualized the study, analyzed data, participated in writing the manuscript.

## Acknowledgments

This work was financed by National Science Centre, Poland through grant number 2016/21/B/NZ9/00840 to MG. The funder had no role in study design, analyzing data and writing the manuscript. We would like to thank Ada Błaszczyk and Anita Kowalczyk for their help in maintaining the plants for the experiments.

---

**Supplementary Table 1.** Osmolytes effect on growth of LB cultures inoculated with lyophilized sugar and sea beet roots untreated with any osmolyte (C), treated with ectoine (E) and trehalose (T). Means of minimum three replicates and standard deviations are given. Statistical significance was assessed with ANOVA and Tukey HSD, significant differences between salinity levels are denoted with different letters.

| NaCl<br>(mM) | *B. huzar* |  |  | *B. maritima* |  |  |
| --- | --- | --- | --- | --- | --- | --- |
|  | C | E | T | C | E | T |
| 0 | 0,395<br>(0,0269) i | 0,445<br>(0,0502) i | 0,472<br>(0,0412) i | 0,338<br>(0,028) fg | 0,528<br>(0,082) f | 0,477<br>(0,0875) f |
| 50 | 0,351<br>(0,0359) h | 0,432<br>(0,0443) h | 0,430<br>(0,0406) h | 0,352<br>(0,051) fg | 0,485<br>(0,125) f | 0,476<br>(0,0537) f |
| 100 | 0,310<br>(0,0450) g | 0,380<br>(0,0482) g | 0,386<br>(0,0322) g | 0,359<br>(0,053) g | 0,403<br>(0,058) e | 0,462<br>(0,0472) f |
| 150 | 0,272<br>(0,0472) f | 0,316<br>(0,0399) f | 0,378<br>(0,0332) f | 0,357<br>(0,029) g | 0,383<br>(0,049) e | 0,438<br>(0,0616) f |
| 200 | 0,194<br>(0,0469) e | 0,235<br>(0,0355) e | 0,267<br>(0,0493) e | 0,312<br>(0,034) f | 0,360<br>(0,051) de | 0,425<br>(0,0752) f |
| 300 | 0,158<br>(0,0418) d | 0,203<br>(0,0277) d | 0,202<br>(0,0359) d | 0,190<br>(0,033) e | 0,306<br>(0,056) cd | 0,322<br>(0,0615) e |
| 400 | 0,078<br>(0,0195) c | 0,115<br>(0,0267) c | 0,154<br>(0,0415) c | 0,164<br>(0,040) de | 0,277<br>(0,055) bc | 0,266<br>(0,0570) d |
| 500 | 0,053<br>(0,0106) bc | 0,099<br>(0,0250) c | 0,132<br>(0,0328) bc | 0,145<br>(0,031) cd | 0,254<br>(0,032) bc | 0,241<br>(0,0542) Cd |
| 600 | 0,041<br>(0,0132) b | 0,059<br>(0,0135) b | 0,109<br>(0,0226) b | 0,125<br>(0,026) bc | 0,226<br>(0,064) b | 0,201<br>(0,0333) bc |
| 700 | 0,004<br>(0,0015) a | 0,007<br>(0,0037) a | 0,007<br>(0,0039) a | 0,102<br>(0,028) ab | 0,157<br>(0,051) a | 0,157<br>(0,0339) ab |
| 800 | 0,004<br>(0,0024) a | 0,006<br>(0,0025) a | 0,007<br>(0,0024) a | 0,078<br>(0,013) a | 0,120<br>(0,040) a | 0,141<br>(0,0297) a |
| 900 | 0,003<br>(0,0021) a | 0,004<br>(0,0029) a | 0,007<br>(0,0029) a | 0,070<br>(0,009) a | 0,102<br>(0,017) a | 0,140<br>(0,0346) a |

